# Transcranial Electrical Stimulation Motor Threshold Combined with Reverse-Calculated Electric Field Modeling Can Determine Individualized tDCS Dosage

**DOI:** 10.1101/798751

**Authors:** Kevin A. Caulfield, Bashar W. Badran, William H. DeVries, Philipp M. Summers, Emma Kofmehl, Xingbao Li, Jeffrey J. Borckardt, Marom Bikson, Mark S. George

**Affiliations:** Brain Stimulation Laboratory, Department of Psychiatry, Medical University of South Carolina, Charleston, SC, USA; Department of Biomedical Engineering, The City College of New York, NY, USA; Ralph H. Johnson VA Medical Center, Charleston, SC, USA

## Abstract

**Background:** Unique amongst brain stimulation tools, transcranial direct current stimulation (tDCS) currently lacks an easy method for individualizing dosage.

**Objective:** Can one individually dose tDCS? We developed a novel method of reverse-calculating electric-field (E-field) models based on Magnetic Resonance Imaging (MRI) scans that can determine individualized tDCS dose. We also sought to develop an MRI-free method of individualizing tDCS dose by measuring transcranial magnetic stimulation (TMS) motor threshold (MT) and single pulse, suprathreshold transcranial electrical stimulation (TES) MT and regressing it against E-field modeling.

**Methods:** In 29 healthy adults, we acquired TMS MT, TES MT, and structural MRI scans with a fiducial marking the motor hotspot. We then computed a “reverse-calculated tDCS dose” of tDCS applied at the scalp needed to cause a 1.00V/m E-field at the cortex. Finally, we examined whether the predicted E-field values correlated with each participant’s measured TMS MT or TES MT.

**Results:** We were able to determine a reverse-calculated tDCS dose for each participant. The Transcranial **<u>Electrical</u>** Stimulation MT, but not the Transcranial **<u>Magnetic</u>** Stimulation MT, significantly correlated with the calculated tDCS dose determined by E-field modeling (R^2^ = 0.509, p < 0.001).

**Conclusions:** Reverse-calculation E-field modeling, alone or in combination with TES MT, shows promise as a method to individualize tDCS dose. The large range of the reverse-calculated tDCS doses between subjects underscores the likely need to individualize tDCS dose. If these results are confirmed in future studies, TES MT may evolve into an inexpensive and quick method to individualize tDCS dose.

## Introduction

Transcranial direct current stimulation (tDCS) is an appealing brain stimulation method due to its efficacy in treating multiple neurological and psychiatric conditions(1–4), relatively cheap cost(5, 6), excellent safety profile(7), and ease of use that could lead to self-administration(7–9). However, tDCS currently does not have a method or biomarker to confirm that stimulation is reaching the cortex or to individualize dose. A typical tDCS study applies a weak uniform electrical current (typically 1-2mA for 20 minutes)(10), often paired with a behavioral task, that may underdose some individuals and be a cause of mixed findings in the field(11–21). Determining a method of individualizing tDCS dosage is important as it would likely inform the experimental design and interpretation of tDCS studies, probably improve the effect size, and allow for more rigorous clinical and investigational use.

Very few studies have examined if there is a way to individualize tDCS dosage. One potential method could be to use electric-field (E-field) modeling combined with a neurophysiological measurement such as transcranial magnetic stimulation (TMS) motor threshold (MT)(22). Researchers have shown that TMS MT correlates with the E-field produced by 1mA of tDCS(22). However, no study has yet explored how to use E-field modeling and a neurophysiological measurement to prospectively individualize tDCS dosage; studies to date have used retrospective tDCS E-field modeling or for only a uniform dose such as 1mA.

To prospectively individualize tDCS dosage using E-field modeling, a first step is to decide upon a desired E-field threshold at the cortex. Currently, there is no consensus about the amount of stimulation it would take to excite cortical tissue using tDCS, likely owing to different neuronal cell types firing at varying thresholds and difficulty with assessing such a threshold in vivo(23). A controversial study by Vöröslakos and colleagues (2018) used implanted electrodes in human cadavers and anesthetized rats to measure the intracortical E-fields produced by tDCS from electrodes on the scalp(24). While many tDCS researchers disagree with the conclusions of the study, Vöröslakos and colleagues determined that an E-field of at least 1mV/mm (equivalent to 1V/m) at the cortex is required to affect neuronal spiking and subthreshold currents; they further estimate that it would likely take 4-6mA of tDCS current to produce E-fields of the 1mV/mm magnitude(24). In this study, we chose to use the 1V/m threshold that was informed by Vöröslakos et al.’s study. An additional benefit of using 1V/m is that it is easily scalable to the desired E-field by multiplication or division. For example, if 0.5V/m is the desired threshold to individualize tDCS dosage to, the calculated tDCS dose to produce 1V/m would be halved.

We aimed to develop a novel method of using E-field modeling to determine an individualized, “reverse-calculated tDCS dose” to produce an E-field threshold of 1.00V/m that could easily be scaled up or down. We correlated the theoretical reverse-calculated tDCS dose with acquired values of TMS MT and transcranial electrical stimulation (TES) MT to determine if a neurophysiological measure could be used to predict individualized reverse-calculated tDCS dose. We hypothesized that TMS MT or TES MT would correlate with reverse-calculated tDCS dose for a 1.00V/m E-field and could be used in the future to individually titrate tDCS dosage to any desired threshold.

## Materials and Methods

### Study Overview

We enrolled 30 healthy participants (15 women, mean age = 26.9, SD = 9.1) in this two visit IRB-approved study. One participant dropped out prior to receiving the MRI scan, so our final sample size was 29. Each participant gave written, informed consent before starting the experimental protocol. In Visit 1, we acquired a resting TMS MT for each participant by stimulating the left motor cortex and recording motor evoked potentials (MEPs) in a standard, closed-loop TMS MT acquisition protocol. We used three electrodes on the contralateral right hand, combined with Spike2 software, to record MEPs. We defined an MEP as having a peak-to-peak amplitude greater or equal to 0.05mV. Parametric Estimation via Sequential Testing (PEST) software was used to help optimally determine the TMS MT in as few pulses as possible(25).

Following TMS MT acquisition, we then acquired an active TES MT for each participant by placing an anodal tab electrode (Natus Neurology, Inc., Pleasanton, CA, USA; rectangular with dimensions of 35 × 20mm) over the TMS motor hotspot and a cathodal ground plate electrode (Natus Neurology, Inc., Pleasanton, CA, USA; rectangular with area = 55 × 42mm) over the left deltoid (**See** **Figure 1**, Supplemental Material S1, and Supplemental Video for a detailed description of the TES procedure). Briefly, participants were instructed to make a “thumbs-up” sign with their contralateral right hands to active the motor circuit, which Merton and colleagues (1982) had previously shown lowers the TES MT by approximately 20%(26). Single, suprathreshold TES pulses were delivered using a constant current stimulator (Digitimer DS7A, Letchworth Garden City, England, UK). With a pulse width of 200μs, a maximum voltage of 400V, monophasic waveform, and an initial stimulation intensity of 58.0mA, we used a modified PEST program to determine the TES MT using only 5 pulses and were able to acquire a relatively painless TES MT for each participant (See Supplemental Material S2 for painfulness and tolerability ratings).

**Figure 1:**
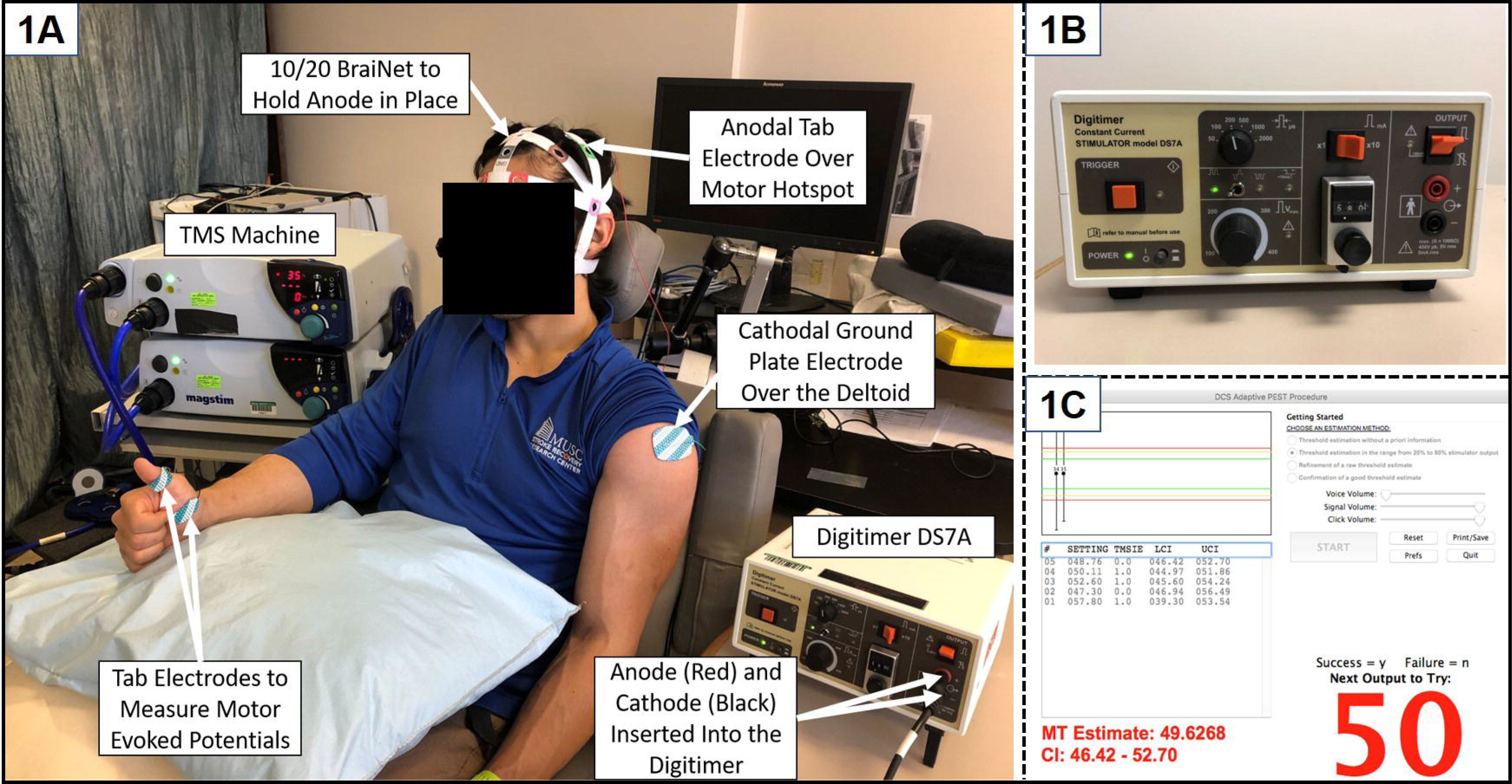
TES Electrode Set-Up. **1A**: Experimental set-up with labeled devices and electrodes. **1B:** A picture of the constant current stimulator (Digitimer DS7A) used to acquire TES MT. **1C:** PEST program window showing an example in which 5 pulses of TES determined a TES MT of 50mA.

In Visit 2, each participant underwent a structural magnetic resonance imaging (MRI) scan. A vitamin E capsule was used as a fiducial to mark each participant’s previously determined scalp target for the TMS/TES MT. This allowed the motor hotspot location to later be visualized in MRICro.

### Preparing MRI Scans for E-Field Modeling

We used the MRICro program to visualize the fiducial marking the left hemispheric motor hotspot location and noted the X, Y, and Z coordinates for the fiducial location (**See** **Figure 2**). In addition, we approximated the cathodal electrode location over the left deltoid by finding the lowest location on the left shoulder visible on the MRI scan (**See** **Figure 1**). For our E-field modeling, we used “Realistic vOlumetric-Approach to Simulate Transcranial Electric Stimulation” (ROAST)” software(27), which allowed us to use the individualized electrode placements, sizes, and orientations used to determine TES MT.

**Figure 2:**
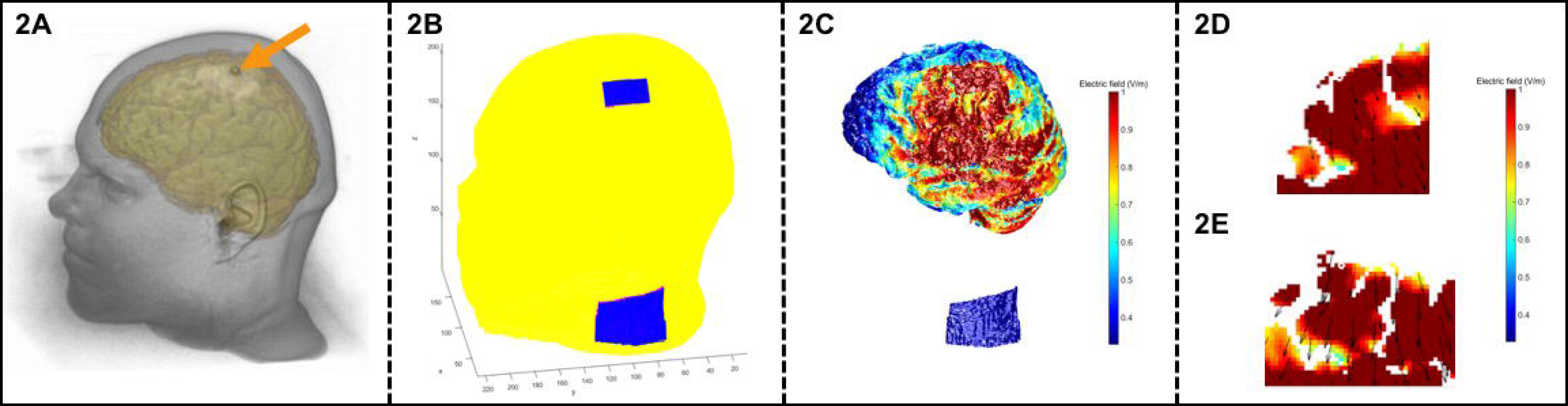
ROAST E-Field Modeling Pipeline for One Participant (1cm Diameter Circular Electrodes). **2A:** Structural MRI with an arrow pointing at the fiducial on the scalp indicating the motor hotspot coordinates (visualized in MRICroGL). **2B:** Using ROAST, an anodal electrode was placed at the left motor hotspot and the cathode was placed on the left shoulder to match the TES electrode montage. **2C:** ROAST E-field model output after skin, skull, CSF, and brain tissue segmentation. **2D/2E:** Close-up views of coronal (2D) and axial (2E) slices with arrows indicating the voxel directly underneath the center of the fiducial marking the motor hotspot. In this example, the E-field was exactly 1.00V/m at this voxel.

### Prospective ROAST E-Field Modeling

Most tDCS E-field modeling studies use modeling to determine the E-field produced by a uniform tDCS current placed on the scalp (e.g. What is the E-field produced by 2mA of stimulation?). In this study, we had the opposite question: To produce an E-field of a 1.00V/m at a certain spot in the cortex, what would be the individualized, reverse-calculated tDCS dose for the electrode on the scalp?

We used ROAST V2.7 for our tDCS E-field modeling as it allowed us to customize the electrode sizes and locations for each participant based on their structural MRI scans(27). We customized the TES pad electrode sizes (Anode: 35mm × 20mm × 3mm; Cathode: 55mm × 42mm × 3mm), locations (left motor hotspot, left deltoid), and orientations (anterior-to-posterior) to reproduce the TES montage in the ROAST code using MATLAB R2015a.

### ROAST E-Field Modeling Methodology-Within Individual Analysis

We sought to determine the reverse-calculated tDCS dose that would be necessary to cause a 1.00V/m E-field at the cortex in each individual by reverse-calculating the ROAST model by computing four E-field models per participant. Using the exact electrode locations used to acquire TES MT, we modeled the E-fields produced from tDCS currents of: 1mA, 3mA, 5mA, and 7mA. We plotted these E-field estimates (in V/m) along the X-axis against the tDCS current input on the Y-axis. We also included the point of 0mA input producing a 0V/m E-field for a total of 5 points in the linear regression.

To determine the E-field produced in each model, we measured the E-field at the voxel directly underneath the center of the anodal electrode that was placed over the left motor hotspot. We calculated a linear regression for this intra-individual model and solved the linear equation for the “reverse-calculated tDCS dose” that would produce exactly a 1.00V/m E-field for that subject at that location. We then computed a fifth ROAST model at this reverse-calculated tDCS dose to confirm that the stimulation input produced the 1.00V/m E-field, and accepted values with a range of 0.99-1.01V/m. All reverse-calculated models produced an E-field value in this range.

Our reverse computation may seem overly elaborate, as theoretically, the electric field is linear with applied current. You should be able to run the model (for a given montage and head) for any current (say 1mA). With multiplication (e.g. no regression) you could then scale the current to produce any desired electric field. This ‘shortcut’ may prove true for future work, and general values. However, the reverse-calculated dose that emerges from the individualized linear model by putting in different current amplitudes is not exactly linear, and we sought in this paper to rigorously test for these assumptions. In the future researchers and clinicians might be able to use one model and scale this up or down as a method of reverse-calculating tDCS dose

### ROAST E-Field Modeling × TMS MT and TES MT Methodology-Group Level Analyses

Following E-field modeling, we plotted each individual’s reverse-calculated tDCS dose against their measured TMS MT and used a group level linear regression to determine the relationship between TMS MT and reverse-calculated tDCS dose (**Figure 3**). We used this same method to then assess the relationship between TES MT and the reverse-calculated tDCS dose in a group level E-Field Model x TES MT regression (**Figures 4**). All statistical analyses were conducted in SPSS 25.0 (Armonk, NY: IBM Corp).

**Figure 3:**
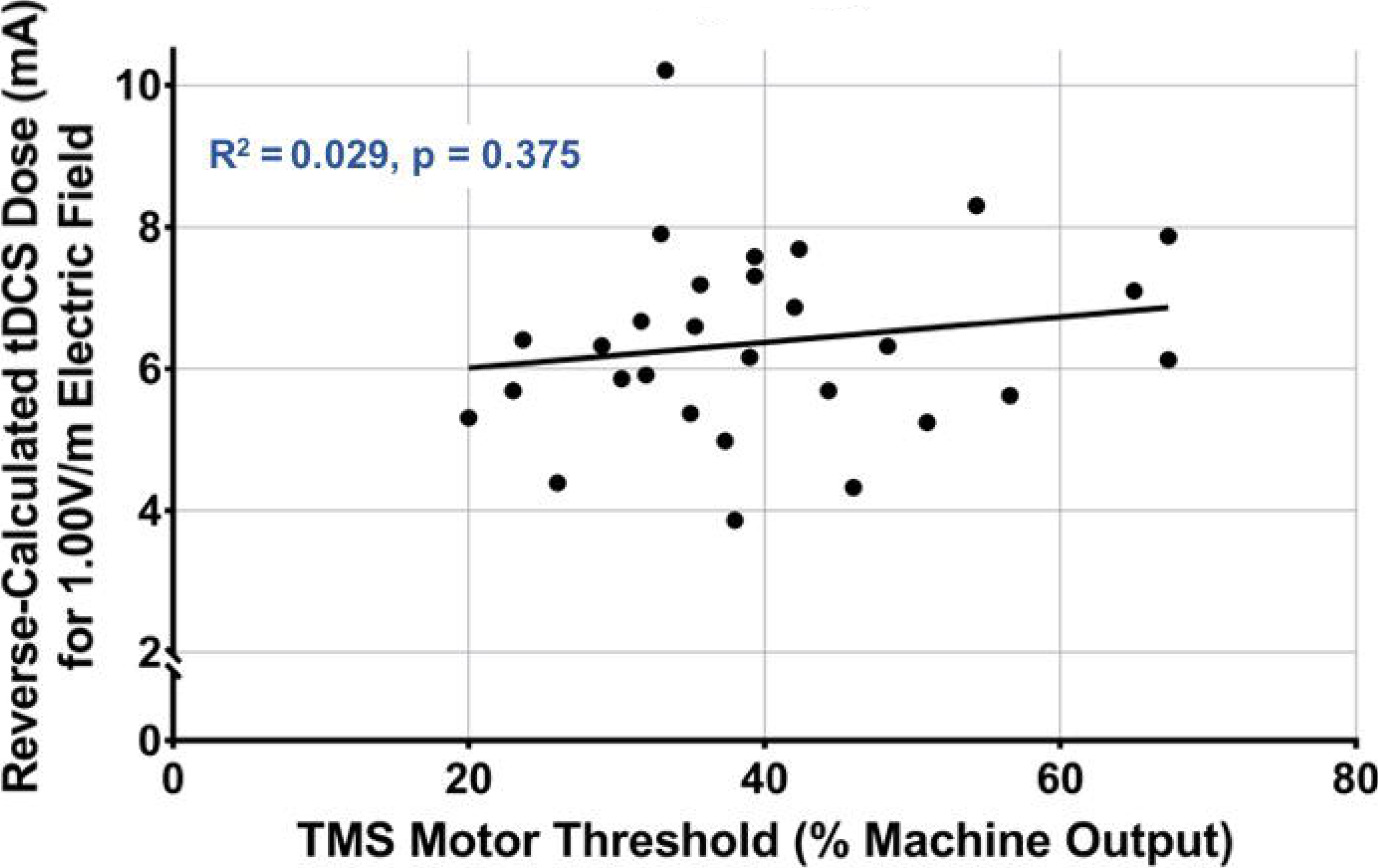
TMS MT Does Not Correlate with Reverse-Calculated tDCS Dose, F(1,27) = 0.813, R^2^ = 0.029, p = 0.375.

**Figure 4:**
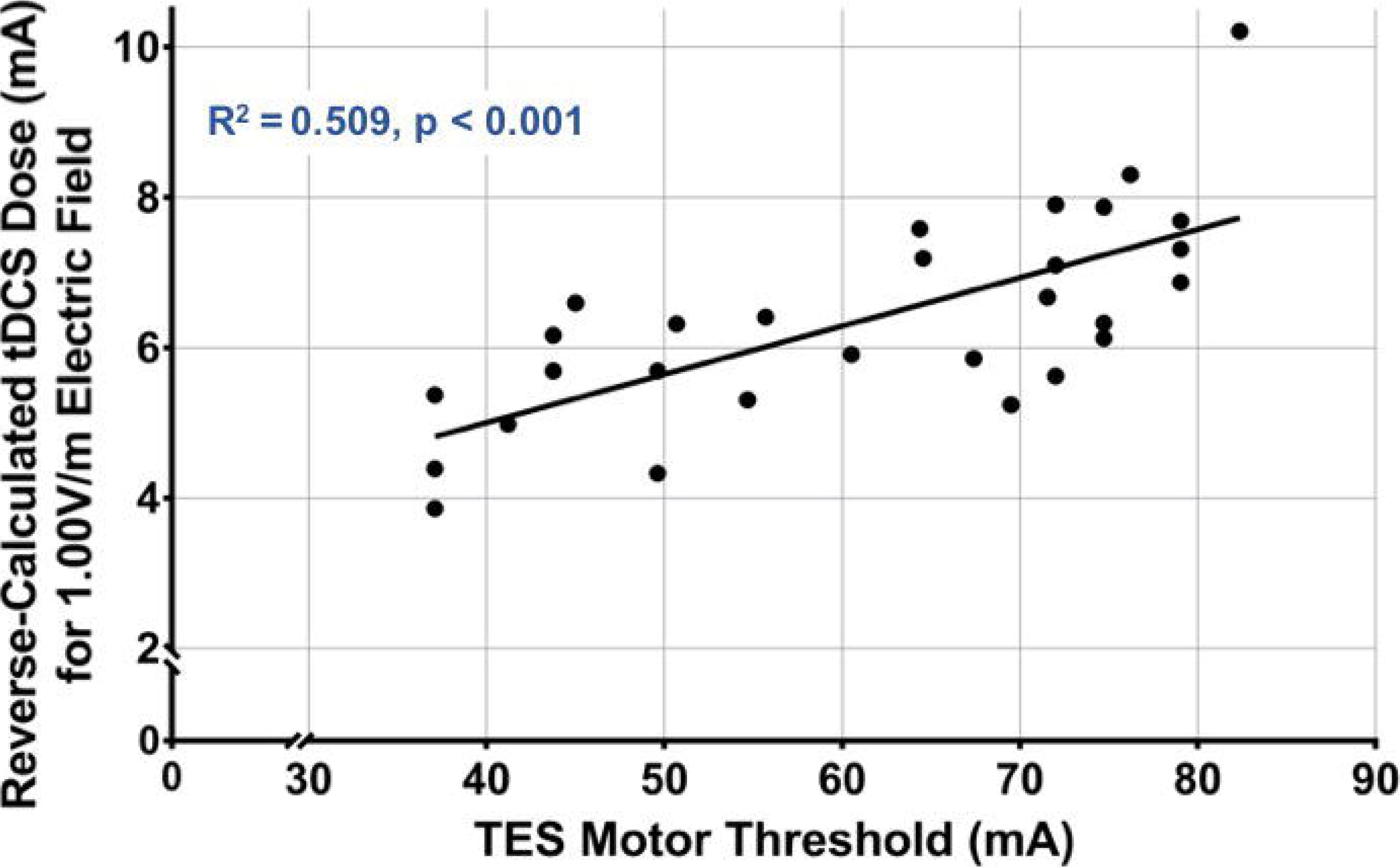
TES MT Significantly Correlates with Reverse-Calculated tDCS Dose, F(1,27) = 27.985, R^2^ = 0.509, p < 0.001.

## Results

### TMS and TES Motor Threshold (MT) Descriptive Statistics

The mean TMS MT was 40.19% of machine output (SD = 12.7%, range = 20-67.3%). The mean TES MT was 61.35mA (SD = 14.91mA, range = 37.1-82.35mA).

### Reverse-calculated tDCS Dose: Actual Electrode Placement and Sizes

The mean reverse-calculated tDCS dose to produce a 1.00V/m E-field in the motor cortex using actual electrode placements was 6.38mA (SD = 1.34mA, range = 3.86 to 10.21mA).

### TMS MT × Reverse-calculated tDCS Dose Linear Regression

This linear regression evaluated the relationship between TMS MT and reverse-calculated tDCS dose determined from the same electrode placement and sizes used for acquiring the TES MT (mean reverse-calculated tDCS dose = 6.38mA, SD = 1.34mA, range = 3.86 to 10.21mA). TMS MT did not statistically predict tDCS dose variance, F(1,27) = 0.813, R^2^ = 0.029, p = 0.375 (**See** **Figure 3**).

### TES MT × Reverse-calculated tDCS Dose Linear Regression

This regression model used the same electrode placement and sizes used to determine the TES MT that were previously used in the TMS MT regression in **Figure 3** (mean reverse-calculated tDCS dose = 6.38mA, SD = 1.34mA, range = 3.86 to 10.21mA). In this regression, TES MT significantly predicted 50.9% of the reverse-calculated tDCS dose variance to produce a 1.00V/m E-field, F(1,27) = 27.985, R^2^ = 0.509, p < 0.001 (**See** **Figure 4**).

The equation for the linear regression is: **Reverse-calculated tDCS Dose = 0.0643 * TES MT + 2.4319**. Thus, measuring a new TES MT and plugging the value into the formula above would allow one to prospectively determine an individual’s reverse-calculated tDCS dose. For example, if an individual had a TES MT of 60mA, the reverse-calculated tDCS dose to produce a 1.00V/m E-field at their motor cortex would be 6.29mA. Notably, this reverse-calculated tDCS dose for a 1.00V/m E-field at the cortex is easily scalable. For example, the reverse-calculated tDCS dose for a 0.50V/m E-field in the same individual would be: 6.29mA * 0.5 = 3.145mA.

### TMS MT × TES MT Linear Regression

We examined the relationship between TMS MT and TES MT by comparing the measured values for each individual in a linear regression. These values did not significantly correlate, F(1,27) = 2.95, R^2^ = 0.099, p = 0.097.

## Discussion

We conducted a study in 29 healthy individuals in which we used both TMS and TES motor thresholds (MT), combined with anatomical neuroimaging and E-field modelling to determine an individualized dosing paradigm for tDCS. This E-field modeling paradigm, was used to determine an individual’s reverse-calculated tDCS dose to produce an E-field of 1.00V/m. We found that an individual’s transcranial ***electrical*** stimulation (TES) MT predicted the reverse-calculated tDCS dose needed to produce a 1.00V/m E-field at the cortex. A linear regression model using the same electrode sizes and positions as our TES MT acquisition predicted 50.9% of the reverse-calculated tDCS dose variance across our sample.

In contrast, a person’s transcranial ***magnetic*** stimulation (TMS) MT did not correlate with the reverse-calculated tDCS dose to produce a 1.00V/m E-field at the cortex. It is unclear why TES MT but not TMS MT correlates with the modelled tDCS dose, but it is likely that the tDCS modeling better captures electrical energy current than that produced by TMS due to differing mechanisms. Our finding that TMS MT did not correlate with TES MT corroborates the idea that TMS MT may not predict reverse-calculated tDCS dose due to a different mechanism (electromagnetic rather than electrical stimulation).

This study suggests several points. First, it is possible to significantly predict approximately 50% of reverse-calculated tDCS dose variance across a relatively young and healthy cohort of participants by combining TES MT and ROAST E-field modeling. While we acquired and analyzed structural MRI scans for each participant in this study, in the future this regression approach could potentially allow TES MT acquisition alone to determine an individual’s reverse-calculated tDCS dose. However, before this regression comparing TES MT and reverse-calculated tDCS dose can be used widely, our results need to be tested for replication and then shown to be valid in some form of a tDCS study measuring behavioral effects.

Second, the reverse-calculated tDCS dose to cause an E-field at a particular threshold at the cortex varies widely between individuals (3.86 to 10.21mA to produce a 1.00V/m E-field at the cortex). This variability in reverse-calculated tDCS dose is substantial. To illustrate the range of dosage, the individual needing the highest reverse-calculated tDCS dose (10.21mA) in our actual electrode position and size model would need a reverse-calculated tDCS dose that is 265% higher than the individual who needed the lowest reverse-calculated tDCS dose (3.86mA). In addition, the inter-individual variance exists regardless of the intended threshold in any region of the brain. For example, in order to produce a 1.00V/m induced electrical field at the motor cortex, the range of tDCS dose needed was from 3.86 to 10.21 mA (average 6.38 mA tDCS dose at scalp). If we moved the entire scale average to instead average 2.0mA at the scalp, the needed individualized range remains 1.21-3.22mA across the sample. If the average dose of 2.0mA were applied uniformly (similarly to how a uniform dose is applied in every extant tDCS study), it would underdose any individual needing above 2.0mA, particularly the person requiring 3.22mA. Taken in sum, our E-field modeling corroborates the idea that individualized tDCS dose is needed for consistent dosing across individuals and studies.

Third, and perhaps controversially, if a 1.00V/m E-field threshold is necessary to cause a spike in neuronal firing, the results from this study support the idea that a uniform 1-2mA tDCS dose is likely insufficient to reach the cortex with a large effect in many participants. While acknowledging that there may actually be some increases in neuronal resting membrane potential at lower than 1.00V/m, our models using this threshold showed that no participant’s reverse-calculated tDCS dose was below 3.86mA and the average reverse-calculated tDCS dose was 6.38mA. tDCS likely has effects at intensities below the 1.00V/m assumption we used, but depending on the reverse-calculated threshold, these results suggest that some, if not many, individuals are underdosed when uniform doses are used.

### Limitations

There were several limitations of this study. Using E-field modeling, even when it has been validated using intracranial recordings, is inherently theoretical. In addition, the method of reverse-calculating tDCS dose can be further refined. While ROAST E-field modeling nicely accounts for many potential co-factors such as scalp-to-cortex distance and tissue conductivity, there may be other factors that influence response to tDCS, even if the stimulation reaches the cortex. Our regression value of R^2^ = 0.509 in **Figure 4** suggests that using TES MT to determine an individualized reverse-calculated tDCS dose for each participant can predict slightly more than 50% of the dose variance. This is a major step forward from accounting for 0% of dose variance in all extant tDCS studies that use uniform doses of current. However, this also means that the source of approximately 50% of the dose variance remains to be determined. A future dose-response study would help to elucidate if individualized dose improves response to tDCS and what other factors may influence reverse-calculated tDCS dose. The relationship between TES MT and reverse-calculated tDCS dose might also change outside of the motor cortex or with different electrode montages (e.g. left M1-supraorbital) and are ongoing areas of research in our lab.

Lastly, it remains unclear what E-field magnitude to target when calculating the reverse-calculated tDCS dose. Based on the existing literature(24) and for ease of scalability to a desired E-field threshold, we reverse-calculated an individualized tDCS dose to produce a 1.00V/m E-field at the cortex for each person. However, it is possible that the 1.00V/m E-field requirement for an increase in neuronal resting potential determined from rodent and human cadaver studies would not scale up to humans or may differ in live human tissue(28). In fact, many tDCS researchers disagree with this 1.00V/m E-field threshold. Thus, using the combination of TES MT and reverse-calculation E-field modeling to individually dose tDCS could potentially be even more informative as the field refines its understanding about the minimally necessary E-field magnitude needed to excite cortical tissue, and then scaling our findings up or down to fit calculate the true reverse-calculated tDCS dosage for each person.

## Conclusions

TES MT is feasible and tolerable. This value, either combined with reverse-calculated E-field modeling or stand alone, can be used to determine a theoretical reverse-calculated tDCS dose for stimulation to reach the cortex of each individual. Our statistical model comparing TES MT to reverse-calculated tDCS dose can be used to individually dose tDCS, predicting approximately 50% of the dose variance in tDCS studies. Moreover, these regressions reveal the wide range (i.e. 3.86 to 10.21mA) between participants, underscoring the need to further develop and evaluate the utility of TES MT combined with E-field modeling for dosing tDCS.

