## Supplemental Materials S1 and S2 for "Transcranial Electrical Stimulation Motor Threshold Combined with Reverse-Calculated Electric Field Modeling Can Determine Individualized tDCS Dosage"

**Supplemental Material S1: Transcranial Magnetic Stimulation (TMS) Motor Threshold (MT) and Transcranial Electrical Stimulation (TES) MT Detailed Methods and Results**

TMS Motor Threshold Protocol

Two rectangular tab electrodes were placed (Natus Neurology, Inc., Pleasanton, CA, USA; electrode area = 35 x 20mm) 0.50cm apart on the subject’s right Abductor Pollicis Brevis (APB) muscle and one ground plate electrode on the back of the hand in a standard procedure for recording motor evoked potentials (MEPs) to determine the resting MT (rMT) (**See Main Text Figure 1A**). Participants wore caps used for marking the brain stimulation target and sat in a chair with a TMS coil holder and fixed head frame. A trained operator then administered single pulses of TMS over the left motor cortex to determine the best location for stimulating the right anterior pollicis brevis (APB) muscle (TMS motor hotspot). A closed-loop TMS-EMG system using Spike2 software combined with a PEST algorithm(1) was used to determine the TMS rMT. Participants then rated the pain, discomfort, and unpleasantness of TMS on a scale from 1 (lowest) to 10 (highest).

TES Motor Threshold Protocol

We marked the center of the TMS coil on the cap before cutting a pinhole to directly mark this spot on the scalp. This location was used as the anodal TES target. We then removed the cap and prepped the scalp target using 70% isopropyl alcohol. One self-adhesive Disposable Tab Electrode (Natus Neurology, Inc., Pleasanton, CA, USA; rectangular with dimensions of 35 x 20mm) was placed on the motor hotspot and connected to the anodal terminal of the constant current stimulator (Digitimer DS7A, Letchworth Garden City, England, UK) **(See Main Text Figures 1A and 1B)**. The anodal tab electrode was oriented with the longer axis running anterior/posterior on the head (**See Main Text Figure 1A**). Next, we prepped the left deltoid using a 70% isopropyl alcohol wipe and placed a self-adhesive Disposable Ground Plate Electrode (Natus Neurology, Inc., Pleasanton, CA, USA; rectangular with area = 55 x 42mm) on this spot as the cathode and inserted this electrode into the Digitimer’s cathodal terminal.

We set the Digitimer’s initial current intensity to 58.0mA (Vmax = 400V, pulse width = 200μs) (**See Main Text Figure 1A).** A modified PEST algorithm and 5 TES pulses over 5 minutes were used to determine the TES active MT (acquired with each participant making the “thumbs up” sign with the contralateral right hand)(1) **(See Main Text Figure 1A and Supplementary Video).** We chose to use an active MT to potentially lower the pain and discomfort felt from each TES pulse. Based on whether or not there was a MEP above 0.05mV in the Spike2 software, PEST determined the next current to try. The TES operator then adjusted the current accordingly (**See Main Text Figure 1C and Supplementary Video**).

**Supplemental Material S2: Statistical Evaluation of Pain, Discomfort, and Unpleasantness Ratings for TMS MT, TES MT; Comparison to 10Hz rTMS Pain Ratings**

TMS Motor Threshold and TES Motor Threshold Overview

For 22 of the 29 participants included in the study, we acquired pain, tolerability, and unpleasantness ratings after both TMS and TES motor thresholding protocols. Following MT acquisition, participants rated either TMS or TES on pain, discomfort, and unpleasantness on scales of 1 (lowest) to 10 (highest).

Statistical Analysis

We compared the pain, discomfort, and unpleasantness ratings of single pulses of TES and TMS using paired t-tests. Next, we used one-sample t-tests to compare our measured TES and TMS pain values to the pain associated with 10Hz rTMS in the OPT-TMS trial for depression, in which 68 patients receiving real rTMS rated the painfulness of 10Hz rTMS on a visual analog scale from 0 to 100(2). We divided the OPT-TMS painfulness scores by 10 to create a range (0 to 10) comparable to that of our TES data (1 to 10). All statistical analyses were conducted in SPSS 25.0 (Armonk, NY: IBM Corp).

Single Pulse TES vs. Single Pulse TMS Pain, Discomfort, and Unpleasantness Ratings

Single pulses of TES (M = 3.24, SEM = 0.39) were significantly more painful than single pulses of TMS (M = 1.57, SEM = 0.19), t(21) = 5.63, p < 0.001 (**See Supplemental Figure 1**). In addition, the discomfort (TES = 5.05, SEM = 0.38 vs. TMS = 2.71, SEM = 0.33), t(21) = 7.0, p < 0.001, and unpleasantness (TES = 5.00, SEM = 0.37 vs. TMS = 2.52, SEM = 0.35), t(21) = 6.24, p < 0.001, ratings were also higher than those for TMS. Thus, even with these modifications, a single TES pulse is generally perceived as more painful than is a single TMS pulse, although they are in the same ballpark.

Single Pulse TES vs. 10Hz rTMS Pain Ratings

Patients in the OPT-TMS trial rated 10Hz TMS as significantly more painful than participants in this study rated single pulses of TES. At the start of the first treatment, pain from 10Hz rTMS (M = 7.34, SEM = 0.35) was significantly higher than was pain from single pulses of TES (M = 3.24, SEM = 0.39), t(21) = 10.57, p < 0.001 (**See Supplemental Figure 1**). We additionally evaluated the painfulness of 10Hz rTMS rated after the last treatment session, when patients have increased tolerance of rTMS pain (lower pain ratings). This analysis also showed significantly higher pain ratings for end of treatment 10Hz rTMS (M = 4.16, SEM = 0.38) than for TES (M = 3.24, SEM = 0.39), t(21) = 3.62, p = 0.002 (**See Supplemental Figure 1**).

**Supplemental Figure Legends**

**Supplemental Figure 1: TES pain ratings compared to single pulse TMS and 10Hz rTMS pain ratings and TES MT vs. TMS MT correlation. 2A:** * Significant at p < 0.01; ** Significant at p < 0.001. Single pulses of TES (M = 3.24, SEM = 0.39) were significantly more painful than single pulses of TMS (M = 1.57, SEM = 0.19). However, single pulses of TES were significantly less painful than 10Hz rTMS in the first (M = 7.37, SEM = 0.35) and last session (M = 4.16, SEM = 0.38) of the OPT-TMS trial for drug-resistant depression.
