## Supplementary figures and images for "Transcranial Electrical Stimulation Motor Threshold Combined with Reverse-Calculated Electric Field Modeling Can Determine Individualized tDCS Dosage"

### Supplemental Figure 1

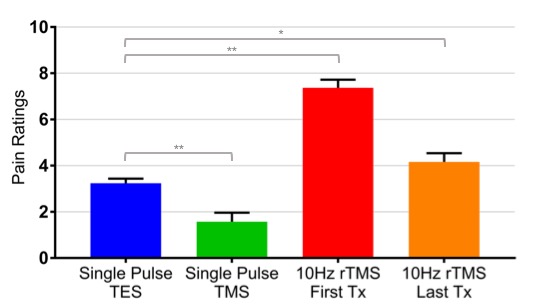
